# Characteristics of the mitochondrial and cellular uptake of MPP^+^, as probed by the fluorescent mimic, 4′I-MPP^+^

**DOI:** 10.1101/321687

**Authors:** Mapa S.T. Mapa, Viet Q. Le, Kandatege Wimalasena

## Abstract

The discovery that 1-methyl-4-phenylpyridinium (MPP^+^) selectively destroys dopaminergic neurons and causes Parkinson’s disease (PD) symptoms in mammals has strengthened the environmental hypothesis of PD. The current model for the dopaminergic toxicity of MPP^+^ is centered on the uptake into dopaminergic neurons, accumulation into the mitochondria, inhibition of the complex-I leading to ATP depletion, increased reactive oxygen species (ROS) production, and apoptotic cell death. However, some aspects of this mechanism and the details of the cellular and mitochondrial accumulation of MPP^+^ are still poorly understood. The aim of this study was to characterize a structural and functional MPP^+^ mimic which is suitable to study the cellular distribution and mitochondrial uptake of MPP^+^ in live cells and use it to identify the molecular details of these processes to advance the understanding of the mechanism of the selective dopaminergic toxicity of MPP^+^. Here we report the characterization of the fluorescent MPP^+^ derivative, 1-methyl-4-(4’-iodophenyl)pyridinium (4’I-MPP^+^), as a suitable candidate for this purpose. Using this novel probe, we show that cytosolic/mitochondrial Ca^2+^ play a critical role through sodium-calcium exchanger (NCX) in the mitochondrial and cellular accumulation of MPP^+^ suggesting for the first time that MPP^+^ and related mitochondrial toxins may also exert their toxic effects through the perturbation of Ca^2+^ homeostasis in dopaminergic cells. We also found that the specific mitochondrial NCX (mNCX) inhibitors protect dopaminergic cells from the MPP^+^ and 4’I-MPP^+^ toxicity, most likely through the inhibition of the mitochondrial uptake, which could potentially be exploited for the development of pharmacological agents to protect the central nervous system (CNS) dopaminergic neurons from PD-causing environmental toxins.

## Introduction

Parkinson’s disease (PD) is characterized by the loss of dopaminergic neurons in the substantia nigra, a region in the midbrain [1, 2]. PD is a chronic and progressive disorder in mid to late ages and characterized by the motor impairment and autonomic dysfunction. The exact cause(s) of dopaminergic neuronal death in PD is not fully understood, but environmental factors are proposed to play a role. The discovery that the synthetic chemical, 1-methyl-4-phenyl-1,2,3,6-tetrahydropyridine (MPTP), recapitulates major pathophysiological characteristics of PD provided the strongest support for the possible environmental contribution to the etiology of PD. Lipophilic MPTP crosses the blood brain barrier and undergoes monoamine oxidase-B catalyzed oxidation in glial cells to produce the terminal toxin, 1-methyl-4-phenylpyridinium (MPP^+^) [3]. Numerous previous *in vivo* and i*n vitro* studies have shown that the metabolite MPP^+^, not the parent compound, MPTP, selectively destroys dopaminergic neurons [4]. Therefore, MPTP/MPP^+^ has been widely used as a convenient model to study the mechanisms of specific dopaminergic cell death in PD and in the development of therapeutic and preventive strategies [5–7].

The currently accepted mechanism for the selective dopaminergic toxicity of MPP^+^ consists several key steps including specific uptake of extracellular MPP^+^ into dopaminergic cells through the plasma membrane dopamine transporter (DAT), active mitochondrial accumulation of cytosolic MPP^+^, inhibition of the complex-I leading to the intracellular ATP depletion, increased reactive oxygen species (ROS) production and apoptotic cell death [8, 9]. Although many aspects of this mechanism have been widely tested and accepted, a number of recent studies have challenged the proposal that the selective in vivo toxicity of MPP^+^ towards dopaminergic cells is due to the specific uptake through DAT, in favor of the possibility that dopaminergic neurons may inherently possess a high propensity towards mitochondrial toxin-mediated ROS production [10,11]. In addition, the molecular details of the mitochondrial accumulation of MPP^+^ is not fully explored or well understood.

Since MPP^+^ is the most widely used model to study the environmental contributions to the etiology of PD at present,[5] a better understanding of the mechanisms of cellular/mitochondrial accumulation and the selective dopaminergic toxicity of MPP^+^ at the molecular level is of importance. Certainly, availability of structural and toxicological MPP^+^ mimics could provide additional information on the cellular distribution, mitochondrial accumulation, and key cellular factors associated with these processes to advance the understanding of the mechanism of the selective dopaminergic cell toxicity of MPP^+^ at the molecular level [12, 13]. In the present study, we have synthesized and characterized 4’I-MPP^+^ as a fluorescent MPP^+^ mimic with desirable toxicological and photophysical properties that could be used to further explore the details of cellular and mitochondrial accumulations of MPP^+^ in live cells to advance the understanding of the mechanism of the selective dopaminergic toxicity of MPP^+^. Using this novel probe, we demonstrate that intracellular Ca^2+^ and the mitochondrial and plasma membrane sodium-calcium exchangers (NCX) play a role in the cellular and mitochondrial accumulation of MPP^+^. Based on these findings, we propose that MPP^+^ and related mitochondrial toxins may also exert their toxic effects through the perturbation of cellular Ca^2+^ homeostasis in dopaminergic cells. In addition, the finding that specific mitochondrial sodium-calcium exchanger (mNCX) inhibitors inhibit the mitochondrial accumulation of MPP^+^ and protect dopaminergic cells from toxicity could potentially be used to develop effective pharmacological agents to protect the central nervous system (CNS) dopaminergic neurons from PD-causing environmental toxins.

## Materials and methods

### Cell lines and material

The mouse hybridoma cell line MN9D [PRID: CVCL_M067[14]] was graciously provided by Dr. Alfred Heller, University of Chicago, IL. Human hepatocellular liver carcinoma cell line [ATCC Cat# HB-8065, RRID: CVCL_0027; (HepG2)] was kind gifted from Dr. Tom Wiese, Fort Hays University, KS. Fresh brains from 12 months old, male, Sprague Dawley rats for mitochondrial isolations were kindly provided by Dr. Li Yao, Wichita State University, KS. The animals used in this study were treated accordance with the recommendations in the Guide for the Care and Use of Laboratory Animals of the National Institutes of Health with the approval of the Wichita State University Institutional Animal Care and Use Committee.

### Chemicals, reagents, and instrumentation

All reagents and supplies were purchased from Fisher Scientific (Pittsburg, PA, USA), Sigma-Aldrich (Milwaukee, WI, USA) or Tocris Bioscience (Bristol, United Kingdom) unless otherwise noted. Krebs-Ringer Buffer-HEPES (KRB-HEPES) contained 109.5 mM NaCl, 5.34 mM KCl, 0.77 mM NaH_2_PO_4_, 1.3 mM CaCl_2_, 0.81 mM MgSO_4_, 5.55 mM dextrose, and 25 mM HEPES at pH 7.4. Dulbecco’s Modified Eagles Medium (DMEM-HCO_3_^-^) contained 109.5 mM NaCl, 5.34 mM KCl, 0.77 mM NaH_2_PO_4_, 1.8 mM CaCl_2_, 0.81 mM MgSO_4_, 44 mM NaHCO_3_, and 5.55 mM dextrose. Freshly isolated rat brain mitochondria were suspended and incubated in media (KCl buffer) containing 0.12 M KCl, 50.0 μM EDTA, 20.0 mM MOPS, pH 7.4

1-methyl-4-(4′-iodophenyl) pyridinium iodide (4’I-MPP^+^) was synthesized using 4-iodophenylmagnisium bromide (prepared from 1-bromo-4-iodobenzene) and N-protected pyridine precursors as previously reported for 4’-halogenated MPP^+^ derivatives [15]. Yield 42 %; mp 305 °C (decomp.);^1^H NMR (d_6_-DMSO)δ 4.3 (s, 3H), 7.8 (d, 2H), 8.0 (d, 2H), 8.5 (d, 2H), 9.0 (d, 2H); ^13^C NMR (d6-DMSO) *δ* 47.0, 100.0, 124.0, 130.0, 138.5, 146.0; MS (ESI) *m*/*z* 295.9804 (M^+^); Fluorescence Ex/Em 320/430 nm. Stock solutions of 4′I-MPP^+^, rotenone, 4′, 6-diamidino-2-phenylindole, dilactate (DAPI), tetramethylrhodamine, methyl ester (TMRM), and 2΄, 7΄-dichlorofluorescin diacetate (DCFH-DA) were prepared in 100 % dimethyl sulfoxide (DMSO). In all experiments, final DMSO concentrations were kept to a minimum usually < 0.05 % v/v.

UV-visible spectra were recorded on a Cary Bio 300 UV-visible spectrophotometer (Varian Inc., Palo Alto, CA). Fluorescence emission spectra were recorded on a Jobin Yvon-Spex Tau-3 spectrophotometer (USA Instruments, Inc., Aurora, OH). The uptake rates of MPP^+^ and 4′IMPP^+^ were determined by HPLC-UV using a solvent system of 48% aqueous buffer containing 20 mM Na_3_PO_4_, 20 mM CH_3_CO_2_Na, 30 mM triethylamine, 1.7 mM 1-octanesulfonic acid sodium salt, pH 7.0, and 52% CH_3_CN at a flow rate of 0.8 mL/min with the UV detection at 295 and 310 nm, respectively. All fluorescence light microscopic experiments were carried out using a Nikon ECLIPS Ti microscope equipped with a Nikon S FLURO 40X objective (Nikon Instrument Inc., Melville, NY). Intracellular ROS levels in live cells were monitored using a Leica [16] confocal fluorescence microscope equipped with a 40X objective (Leica Microsystems Inc., Buffalo Grove, IL).

### Methods

### Cell culture

MN9D [14] and HepG2 [17] cells were cultured in 100 mm^2^ Falcon tissue culture plates in DMEM-HCO_3_^-^ supplemented with 10% fetal bovine serum, 50 μg/mL streptomycin, and 50 IU/mL penicillin at 37 °C in a humidified atmosphere of 7% CO_2_. Cells were cultured to about 70–80% confluence and then seeded into glass-bottomed culture plates (for imaging studies) or 12- or 96-well plates depending on the experiment and grown to about 70–80% confluence (usually two days) unless otherwise stated.

### Differentiation of MN9D cells

The differentiation of MN9D cells was carried out with 1.0 mM sodium *n*-butyrate according to the published procedure of Choi *et al* [14]. Briefly, MN9D cells were grown in DMEM-HCO_3_^-^ containing 10% fetal bovine serum with 1.0 mM sodium butyrate at 37 °C in a humidified atmosphere of air with 7% CO_2_ for 3 days at which time the media was replaced with fresh media containing 1.0 mM sodium *n*-butyrate and incubated for an additional 3 days. The progress of differentiation was judged by observing the expected morphological changes. All differentiated cells were used in appropriate experiments after the 6^th^ day of differentiation.

### Time dependency of cellular uptakes of MPP^+^ and 4′I-MPP^+^

Cells were seeded into 12-well plates at 0.25 × 10^6^ cells/well and grown 2–3 days to about 70–80% confluency. The media was removed and a solution containing 100 μM MPP^+^ or 4′I-MPP^+^ in warm KRB-HEPES (or without Ca^2+^ and with 1.4 mM EGTA) and was added to each well and incubated at 37 °C for the desired period of time (*see Figure legends for further details). After the* incubations, the media were removed and cells were washed with ice-cold KRB-HEPES. Washed cells were suspended in 1.0 mL of ice-cold KRB-HEPES and 50 μL aliquots were withdrawn for protein determination. The remaining cell suspensions were centrifuged at 5000 × g at 4 °C for 5 min and the cell pellets were treated with 75 μL of 0.1 M HClO_4_. The coagulated proteins were pelleted by centrifugation at 13,200 × *g* at 4°C for 8 min and the MPP^+^ or 4′I-MPP^+^ contents of the supernatants were determined by C_18_-reversed-phase HPLC-UV (HPLC-UV) as previously described [18] using standard curves constructed from authentic standards. The standard curves for HPLC-UV quantifications were constructed using authentic samples of MPP^+^ and 4′I-MPP^+^ under the same HPLC-UV separation conditions (S1 Fig). All MPP^+^ and 4′I-MPP^+^ levels were normalized to the respective protein concentration of each sample and were corrected for non-specific membrane binding by subtracting the corresponding time zero readings for each MPP^+^ and 4′I-MPP^+^ concentration (normally <1% of the intra-cellular concentration).

To quantify 4′I-MPP^+^ uptake by fluorescence, MN9D or HepG2 cells grown in glass bottomed plates were washed, reconstituted with KRB-HEPES (1.0 mL), pH 7.4, and mounted on the stage of Nikon ECLIPSE Ti-S inverted fluorescence microscope. After the ROIs were selected, cells were incubated for 2 min and 4′I-MPP^+^ was added to a final concentration of 100 μM and the intracellular fluorescence at Ex/Em 340/470–550 nm (although the excitation maxima of 4′I-MPP^+^ was 320 nm, we used excitation wavelength of 340 nm to minimize the exposure of cells to UV) was recorded as a function of time for period of 1 h. The averages of background corrected, control subtracted ROI fluorescence intensities were used to estimate the cellular uptake of 4′I-MPP^+^.

### Measurement of cell viabilities

Cell viability was determined by the MTT [3-(4,5-dimethylthiazol-2-yl)-2,5-diphenyltetrazolium bromide)] assay [19]. Briefly, cells were seeded on 96-well plates and allowed to grow to about 70–80% confluence. Prior to the experiment, culture media was replaced with DMEM-HCO_3_^-^ containing desired concentrations of MPP^+^ or 4′I-MPP^+^ and incubated for 16 h at 37 °C. After the incubation, 10 μL of 5 mg/mL MTT solution was added to each well and incubated for 2 h at 37 °C. The resulting formazan was solubilized by the addition of 210 μL detergent solution [50% DMF/H_2_O (v/v), 20% SDS (w/v)] followed by incubation at 37 °C for 12 h. The cell viabilities were determined by quantifying solubilized formazan by measuring the difference in the absorbance at 570 nm and 650 nm [20]. Results were expressed as % viability of MPP^+^ or 4′I-MPP^+^ treated cells with respect to control cells, which were treated under the same conditions except in the absence of the MPP^+^ or 4′I-MPP^+^.

### Measurement of mitochondrial complex-I inhibition

Isolated rat brain mitochondria were lysed by freeze-thawing in hypotonic media containing 25 mM K_3_PO_4_ and 5 mM MgCl_2_, pH 7.2 [21]. The complex-I (i.e. NADH: ubiquinone oxidoreductase) activity of mitochondrial membrane fragments were determined according to the procedure of Birch-Machin and Turnbull [22]. Briefly, mitochondrial membranes (50 μg protein) were incubated with 130 μM NADH (final concentration) and Antimycine A (2 μg/mL) in a 1.0 mL assay solution containing 25 mM K_3_PO_4_, 5 mM MgCl_2_, 2 mM KCN, 2.5 mg/mL bovine serum albumin, pH 7.2, for 2 min at 37°C with or without rotenone, MPP^+^ or 4′I-MPP^+^ (for further details see the corresponding figure legends). The complex-I mediated ubiquinone dependent NADH oxidation was initiated by adding ubiquinone-1 to a final concentration of 65 μM and the initial rates of the reactions were monitored by following the decrease in absorbance at 340 nm with respect to 425 nm reference wavelength for 5 min.

### Measurement of intracellular ATP levels

Cells grown in 12-well plates were treated with 100 μM MPP^+^ or 4′I-MPP^+^ in KRB-HEPES for 6 h at 37 °C. After the treatments, cells were suspended in 1 mL of ice-cold KRB-HEPES and 50 μL samples were withdrawn for protein quantification. The remaining cell suspensions were centrifuged for 3 min at 3,500 × g at 4 °C and the cell pellets were treated with 1 mL of lysis buffer (0.1 M Tris, 1% Triton X-100, pH 7.5) for 2 min and centrifuged at 3,500 × g for 3 min to remove the cell debris. The ATP contents of the supernatants were measured using a fluorometric ATP assay kit (BioVision, Milpitas CA, USA) following the manufactures guidelines. All ATP levels were normalized to the protein content of each sample.

### Measurement of intracellular Reactive Oxygen Species (ROS)

Cells grown in 12-well plates were incubated with 10 μM DCFH-DA in KRB-HEPES for 1 h. DCFH-DA-loaded cells were washed with ice-cold KRB-HEPES and treated with 100 μM MPP^+^ or 4′I-MPP^+^ in the same buffer for 1 h at 37°C. After the incubations, cells were washed, harvested, solubilized with 0.1 M Tris buffer (pH 7.5) containing 1% Triton X-100, and cell debris were removed by centrifugation at 13,200 × g for 8 min at 4°C. The content of ROS-oxidized DCFH product, 2΄,7΄-dichlorofluorescein (DCF), in the supernatants were quantified by fluorescence (Ex/Em 504/525 nm) [16]. **The data were normalized to the protein contents of individual samples**. ROS productions in DCFH-DA (50 μM) loaded 250 μM 4′I-MPP^+^ treated (4 h) live MN9D cells were visualized by confocal fluorescence microscopy at Ex/Em 488/524 nm using a Leica TCS SP5II microscope.

### Chromatin condensation test for apoptosis

MN9D cells grown in glass-bottomed plates were treated with 250 μM MPP^+^ or 4′I- MPP^+^ for 12 h and then with 300 nM DAPI in KRB-HEPES for 15 min in the dark at 37 °C [23]. Then cells were rinsed and PBS was added to cover the cell layer. Increase in nuclear DAPI fluorescence (Ex/Em at 358/461 nm) due to chromatin condensation was observed using a Nikon ECLIPSE Ti-S inverted fluorescence microscope. The controls were treated similarly except that MPP^+^ or 4′I-MPP^+^ was omitted from the initial incubation medium.

### Mitochondrial localization of intracellular 4′I-MPP^+^

Cells were grown in glass-bottomed plates and incubated with 100 μM 4′I-MPP^+^ and 200 nM MitoTracker Green FM (Life Technologies, Grand Island NY, USA) in KRB-HEPES at 37 °C for 30 min. Then, cells were washed, reconstituted with KRB-HEPES (1.0 mL), pH 7.4, and dual fluorescence of intracellular 4′I- MPP^+^ (Ex/Em 340/470–550 nm) and MitoTracker Green FM (Ex/Em 644/665 nm) were recorded using a Nikon ECLIPSE Ti-S inverted fluorescence microscope fitted with a 40X objective.

### Measurement of MPP^+^ and 4′I-MPP^+^ uptake into isolate MN9D cell mitochondria

Intact mitochondria were isolated from MN9D cells according to the published procedure of Frezza et al [24]. The isolated intact mitochondrial preparations were suspended in KCl buffer and aliquots (5 μL) were withdrawn for protein determination. Desired amounts of mitochondrial suspensions were incubated with 400 μM MPP^+^ or 4′I-MPP^+^ in KCl buffer for 45 min at 37 °C. After the incubation, incubates were centrifuged at 7,000 × *g* at 4 °C for 10 min and the mitochondrial pellets were treated with 75 μL of 0.1 M HClO_4_. The coagulated proteins were removed by centrifugation at 13,200 × *g* for 8 min at 4°C and MPP^+^ and 4′I-MPP^+^ contents in HClO_4_ extracts were quantified by HPLC-UV as detailed above. All MPP^+^ and 4′I-MPP^+^ levels were normalized to the respective protein concentration of each sample and were corrected for non-specific membrane binding by subtracting the corresponding time zero readings for each MPP^+^ and 4′I-MPP^+^ concentration (normally <1.5% of the intra-cellular concentration).

### Effects of MPP^+^ and 4′I-MPP^+^ on the mitochondrial membrane potentials

MN9D or HepG2 cells grown in glass-bottomed plates were treated with 50 nM TMRM (AnaSpec, Fremont CA, USA) in KRB-HEPES for 45 min in the dark [25]. After mounting on the stage of Nikon ECLIPSE Ti-S inverted fluorescence microscope, 15–20 ROIs with uniform fluorescence were selected and TMRM fluorescence at Ex/Em at 543/573 nm was recorded at 5 sec time intervals for 3 min. Then, MPP^+^ or 4′I-MPP^+^ was added to a final concentration of 500 or 100 μM, respectively and TMRM fluorescence measurements were continued for additional 40 min for MPP^+^ and 15 min for 4′I-MPP^+^ at 15 sec time intervals. The background fluorescence was subtracted from the ROI fluorescence and averages of corrected data were used to estimate the magnitude of the mitochondrial membrane potential.

### Effect of FCCP on the cellular uptake of MPP^+^ and 4′I-MPP^+^

Cells grown in 12-well plates were washed and incubated with 5 μM TFCCP in KRB-HEPES for 30 min followed by 50 μM MPP^+^ or 4′I-MPP^+^ in the same buffer for 45 min at 37 °C. After the incubations, cells were washed with ice-cold KRB-HEPES and treated with 0.1 M HClO_4_. The intracellular MPP^+^ or 4′I-MPP^+^ contents of HClO_4_ extracts were quantified by HPLC-UV as detailed above. The controls were treated under the same conditions except that FCCP was omitted from the initial incubation. All uptake data were normalized to the respective protein concentration of each sample. To visualize the effect of FCCP on the mitochondrial accumulation of 4′I-MPP^+^, cells were grown in glass bottomed plates and incubated with 5 μM FCCP in KRB-HEPES for 30 min followed by 100 μM 4′I-MPP^+^ for 1 h at 37 °C. Then cells were washed and reconstituted with 1 mL of KRB-HEPES and the intracellular 4′I-MPP^+^ fluorescence (Ex/Em 340/470–550 nm) were recorded.

### The effects of Ca^2+^ and mNCX inhibitors on the MN9D mitochondrial uptake of 4′I-MPP^+^

Intact MN9D cell mitochondria were incubated with 200 μM 4′I-MPP^+^ for 45 min at 37 °C in 0, 500 or 1000 nM Ca^2+^ containing KCl buffer. When mNCX inhibitors (KBR7943, 2APB and CGP37157) were used, mitochondrial suspensions were initially incubated with 10, 25 or 50 μM inhibitor in KCl buffer for 10 min at 37 °C and then with 200 μM 4′I- MPP^+^ for 45 min at 37 °C. After the incubations, samples were centrifuged at 7,000 × *g* at 4 °C for 10 min and the mitochondrial pellets were treated with 75 μL of 0.1 M HClO_4_. The coagulated proteins were pelleted by centrifugation at 13,200 × *g* for 8 min at 4°C and the 4′I-MPP^+^ contents in HClO_4_ extracts were quantified by HPLC-UV as detailed above. To visualize the effect of CGP37157 on the mitochondrial uptake of 4′I-MPP^+^ in live cells, HepG2 cells grown in glass-bottomed plates were incubated with 50 μM CGP37157 for 30 min initially and then with 100 μM 4′I-MPP^+^ and 200 nM MitoTracker Green FM in KRB-HEPES for 1 h at 37 °C. After incubation cells were washed and reconstituted with KRB-HEPES, pH 7.4 and 4′I-MPP^+^ and MitoTracker Green FM fluorescence images of cells were recorded as detailed in above. Controls were treated under the same experimental conditions, except that CGP37157 was omitted from the initial incubation.

### Effects of CGP37157 and KBR7943 on the 4′I-MPP^+^–mediated mitochondrial membrane depolarization

HepG2 cells grown in glass-bottomed plates for two days were treated with 50 μM CGP37157 or KBR7943 in KRB-HEPES for 30 min followed by 50 nM TMRM in KRB-HEPES for 45 min. Then plates were mounted on the stage of the microscope and mitochondrial membrane depolarization upon the treatment with 100 μM 4′I-MPP^+^ was measured as a function of time as detailed above. The effect of CGP37157 on the 4′I-MPP^+^ mediated mitochondrial membrane depolarization at longer time periods was monitored by observing the reduction of TMRM fluorescence from 0 to 4 h incubations in the presence or absence of CGP37157. HepG2 cells were incubated with or without 10 μM CGP37157 followed by 50 nM TMRM, and finally with 4′IMPP^+^ (100 μM, 0 or 4 h) and the intracellular TMRM fluorescence was recorded. Controls were carried out using an identical protocol except that either CGP37157 or 4′I-MPP^+^ were omitted from the incubation media.

### Protein determination

Protein contents of various cell preparations were determined by the method of Bradford using bovine serum albumin (BSA) as the standard [26]. Aliquots of (50 μL) of cell suspensions were incubated with Bradford protein reagents (950 μL) for 10 min and absorbance at 595 nm were recorded. The absorbance readings were converted to the corresponding protein concentrations using a standard curve constructed employing BSA.

### Statistical analyses

All data are averages of three or more replicates of a single experiment and presented as mean ± SD (all the experiments were carried out multiple times and the representative data from a single experiment are shown). To correct for the minor variations in color development in the MTT assay of cell viability, all the absorbance values were converted to percentages of controls, which were treated under the same conditions, except that MPP^+^ and 4′I- MPP^+^ were excluded from the incubations. All quantitative data i.e. uptake, ATP, and ROS levels were normalized to the protein content of individual samples to correct for the cell density variations. P values for pair-wise comparisons were calculated by two-tailed Student’s T-test. For all the data a level of p<0.05 was considered statistically significant.

### Technical statement

Due to the high toxicity and obvious health hazards of MPP^+^ and 4′I- MPP^+^, extreme caution was used in their handling in accordance with published procedures [27].

## Results

### Photophysical properties of 4′I-MPP^+^

4′I-MPP^+^ (Fig 1A) showed a characteristic UV-Vis spectral analysis of absorption band at ∼313 nm and a moderately intense fluorescence band with the excitation and emission maxima at 320 and 430 mm in PBS, respectively (Fig 1B and 1C). Standard photochemical studies showed that neither the absorption band at 313 nm nor the fluorescence emission at 430 nm of 4′I-MPP^+^ was significantly affected upon changing the polarity of the solvent i.e. from acetonitrile to PBS (Fig 1B and 1C; the effect of less polar solvents such as CHCl_3_ could not be determined due to the low solubility). Similarly, the fluorescence of 4′I-MPP^+^ in acetonitrile was quenched by about ∼60% in PBS (Fig 1C). Live cell fluorescence images of 4′I-MPP^+^ (50 μM; Ex/Em 340/470–550 nm) treated differentiated MN9D cells show that intracellular 4′I-MPP^+^ could be easily detected by its intrinsic fluorescence (Fig 1D). Data in Fig 1E show that 4′I-MPP^+^ uptake time courses determined independently by HPLC-UV [HPLC-UV quantifications of intracellular MPP^+^ and 4′I-MPP^+^ levels using standard curves constructed from authentic standards of MPP^+^ and 4′I-MPP^+^ (S1 Fig.)] or fluorescence (Ex/Em 320/430 nm) measurements were similar for both MN9D and HepG2 cells. The results further show that MN9D cells uptake time courses of both toxins (at 100 μM concentration) were linear during the 60 min incubation period. However, the rate of 4′I-MPP^+^ uptake was about twice as fast as MPP^+^ (1.2 nmoles/mg versus 0.6 nmoles/mg of protein in 1 h; Fig 1E). The rates of uptake of both MPP^+^ and 4′I- MPP^+^ into HepG2 cells were apparently saturable and relatively faster than that of MN9D cells under similar experimental conditions (Fig 1E). In addition, while HepG2 MPP^+^ uptake reached to about 1.2 nmoles/mg in 60 min, 4′I-MPP^+^ uptake reached to 1.4 nmoles/mg of protein in 40 min, again suggesting that the rate of 4′I- MPP^+^ uptake is faster than that of MPP^+^, parallel to that observed for MN9D cells. Further studies revealed that the non-specific membrane binding of 4′I-MPP^+^ (or various membrane transporters) was relatively small [<1% under the standard experimental conditions (data not shown).

**Fig 1.**
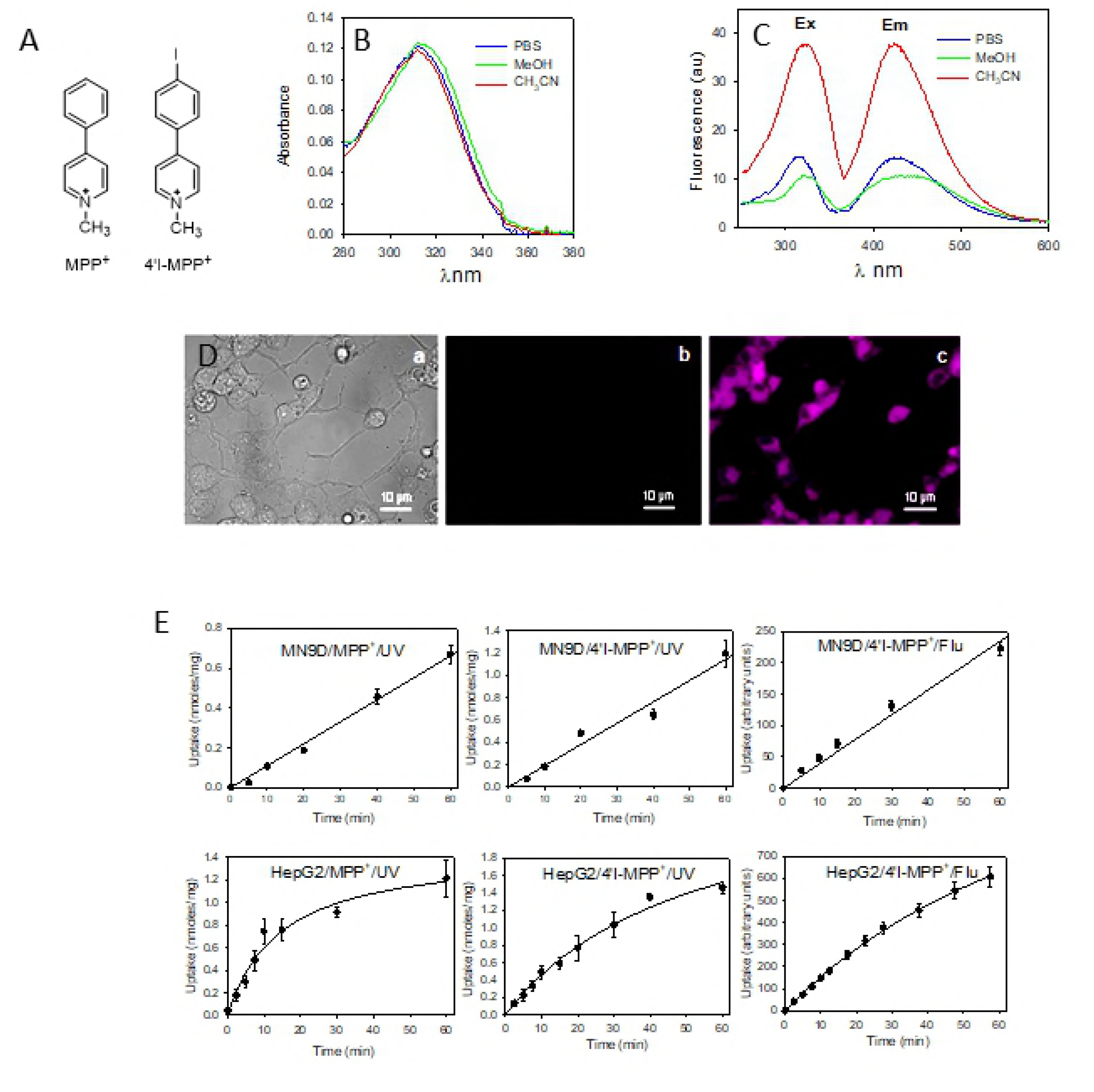
Structure, photophysical properties and intracellular accumulation of 4′I-MPP^+^. (A) Structures of MPP^+^ and 4′I-MPP^+^; (B) UV-Vis spectra of 4′I-MPP^+^ (5 μM) in PBS, methanol, and acetonitrile; (C) Excitation and emission spectra of 4′I-MPP^+^ (5 μM) in PBS, methanol, and acetonitrile); (D) Light and Fluorescence (Ex/Em 340/470–550 nm) images of 4′I-MPP^+^ (50 μM) treated differentiated MN9D cells. (a) light image; (b) florescence image at zero time after 4′IMPP^+^ treatment; (c) florescence image after 2 h after 4′I-MPP^+^ treatment of the same cell population; (E) The time courses of the cellular uptakes of 100 μM MPP^+^ and 4′I-MPP^+^ which were determined either by HPLC-UV [15] or by measuring the intrinsic fluorescence of intracellular 4′I-MPP^+^.

### Relative cell toxicities of MPP^+^ and 4′I-MPP^+^

The selective dopaminergic M9ND cell toxicity of 4′I-MPP^+^ was determined relative to MPP^+^ employing HepG2 cells as a non-dopaminergic control, using the MTT cell viability assay. The data show that while MN9D cells are highly susceptible, HepG2 cells are resistant to both toxins during 16 h incubation periods, under similar experimental conditions (Fig 2A). The data further show that the IC_50_ for MPP^+^ is about 125 μM, while that for 4′I-MPP^+^ is about 40 O Ȉ suggesting 4′I-MPP^+^ is relatively more toxic to MN9D cells, in comparison to MPP^+^.

**Fig 2.**
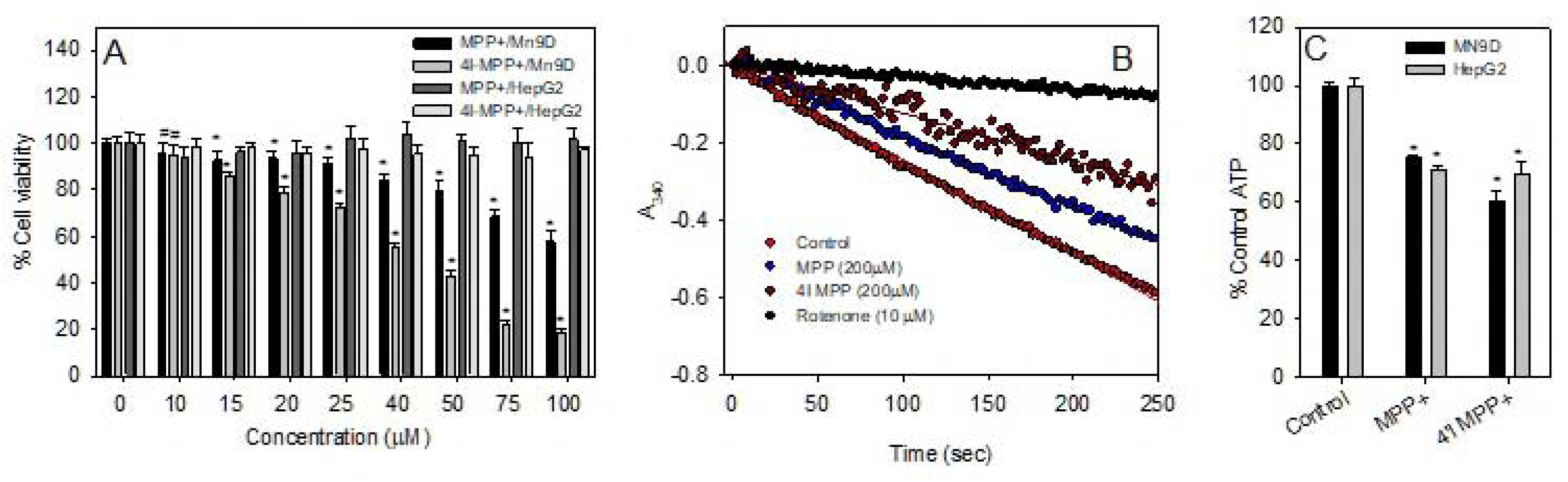
Comparison of the toxicological characteristics of 4′I-MPP^+^ with respect to MPP^+^. (A) 4′I-MPP^+^ is selectively toxic to dopaminergic MN9D (reference to HepG2) cells similar to MPP^+^ as determined by MTT assay. Data are presented as mean ± SD (n = 6). ^*^p<0.005 and #p<0.05 versus no toxin controls; (B) Both MPP^+^ and 4′I-MPP^+^ (200 μM) inhibit the ubiquinone dependent NADH oxidation activity of rat brain mitochondrial complex-I. The complex-I inhibition potency of 4′I-MPP^+^ is greater than that of MPP^+^ at 200 μM concentrations; (C) Both MPP^+^ and 4′I-MPP^+^ deplete intracellular ATP levels. Intracellular ATP levels of MPP^+^ or 4′I-MPP^+^ (100 μM 8 treated MN9D and HepG2 cells (for 6 h at 37 °C) were quantified as detailed Materials and methods; The data presented as mean ± S.D (n = 3). ^*^p< 0.0025 versus the controls with no toxin.

### 4′I-MPP^+^ and MPP^+^ causes mitochondrial complex-I Inhibition, ATP depletion, increased intracellular ROS production causing apoptotic cell death in dopaminergic cells

The effect of 4′I-MPP^+^ on the NADH-ubiquinone oxidoreductase (complex-I) activity of rat brain mitochondrial membrane fragments was determined using the ubiquinone dependent NADH- oxidation assay [22]. The inhibition of complex-I was measured by measuring the decrease in absorbance due to the ubiquinone dependent oxidation of NADH at 340 nm as a function of time. Under the standard assay conditions, rotenone, a well-characterized specific and potent complex-I inhibitor [28] inhibited about 90% of the NADH oxidation activity at 10 μM concentration confirming the suitability of the assay for specific complex-I activity measurements (Fig 2B). Under similar conditions, 4′I-MPP^+^ and MPP^+^ showed 46% and 21% inhibition of the complex-I activity at 200 μM concentrations, respectively, suggesting that 4′I-MPP^+^ is a better complex-I inhibitor than MPP^+^ (Fig 2B). In addition, both MPP^+^ and 4′I-MPP^+^ deplete intracellular ATP levels in both MN9D and HepG2 cells, with 4′I-MPP^+^ showing a more pronounced effect in comparison to MPP^+^ in MN9D cells (Fig 2C). Furthermore, both MPP^+^ and 4′I-MPP^+^ treatments increase ROS production [measured by the ROS sensitive fluorescent probe DCFH-DA [29]] specifically in MN9D (Fig 3A), but not in HepG2 cells. Fluorescence live cell imaging experiments also show the 4′I-MPP^+^–mediated intracellular ROS production in MN9D (Fig 3B). 4′I-MPP^+^–mediated MN9D apoptotic cell death is associated with the characteristic apoptotic cell death pathway including cell shrinkage and chromatin condensation (as indicated by the increase of the nuclear DAPI fluorescence in toxin treated cells), similar to the MPP^+^ (Fig 3C).

**Fig 3.**
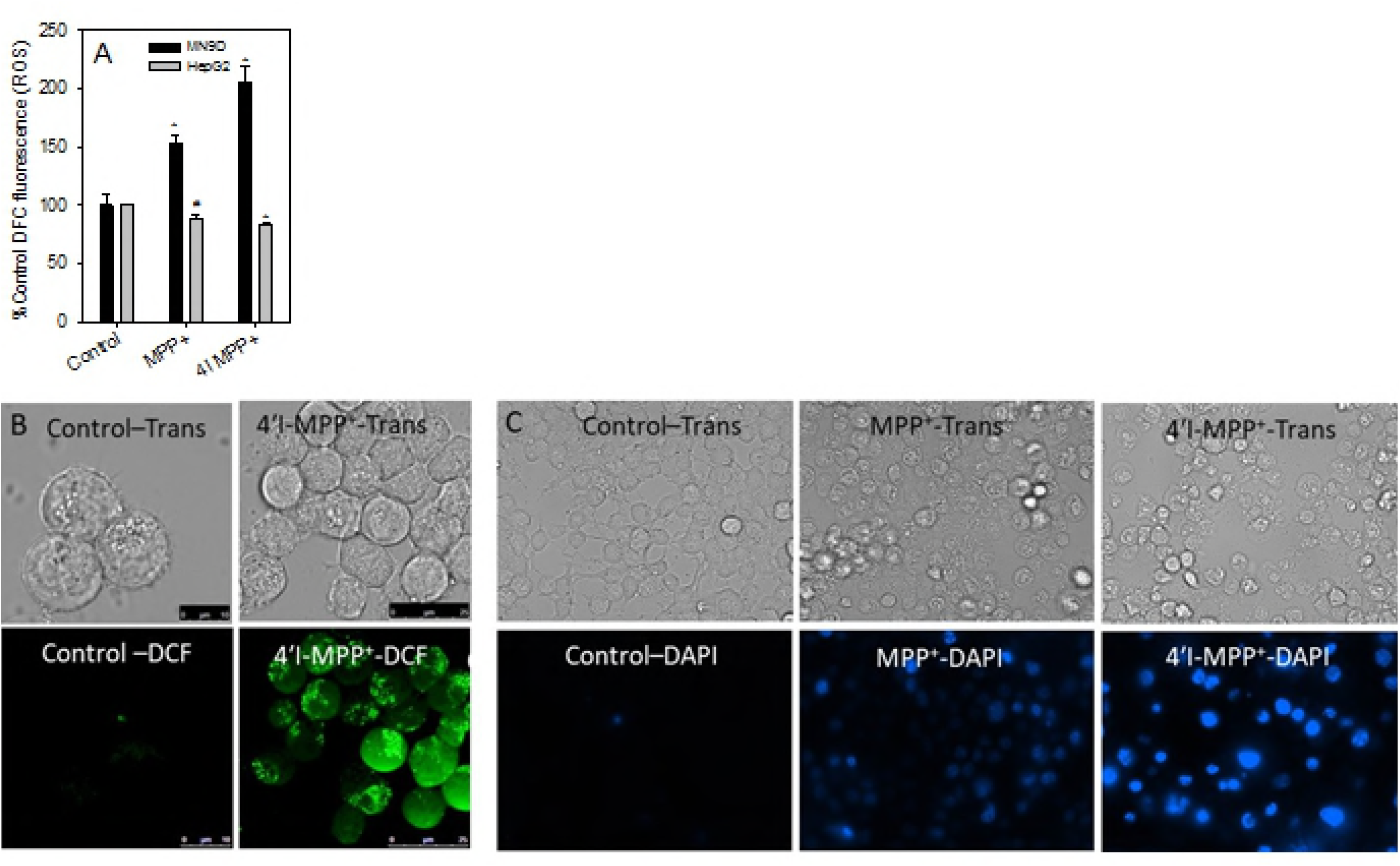
Increase of intracellular ROS levels by 4′I-MPP^+^ and MPP^+^ leading to specific apoptotic cell death. (A) The intracellular ROS levels were determined using the non-specific ROS sensitive probe DCFH-DA in MPP^+^ or 4′I-MPP^+^ treated (100 μM, 1 h at 37 °C) MN9D and HepG2 cells as detailed in Materials and methods. The data represent as mean ± S.D (n = 3). ^*^p<0.005 and #p<0.02 compared with controls with no toxins; (B) 4′I-MPP^+^–mediated ROS production in MN9D cells is localized to the mitochondria. The DCF fluorescence (Ex/Em 488/524 nm) images of 4′I-MPP^+^ treated (250 μM for 4 h at 37 °C) MN9D cells were recorded as detailed in Materials and methods; (C) Both MPP^+^ and 4′I-MPP^+^ cause apoptotic MN9D cell death. Apoptotic chromatin condensation was visualized by observing the increase in nuclear DAPI fluorescence (Ex/Em 358/461 nm) in MPP^+^ or 4′I-MPP^+^ (250 μM treated (for 12 h) MN9D cells.

### Mitochondrial uptake and intracellular localization of 4′I-MPP^+^

The intracellular localization of 4′I-MPP^+^ in differentiated and undifferentiated MN9D and HepG2 cells was examined using the mitochondrial marker, MitoTracker Green FM. As shown from the data in Fig 4A, the fluorescence of MitoTracker Green FM overlaps well with bulk of the intracellular 4′I-MPP^+^ fluorescence in all cases, suggesting that intracellular 4′I-MPP^+^ is primarily localized into the mitochondria of both cell types without significant cell-type specificity. The fluorescence images further show that no significant fluorescence is associated with the cell nucleoli or the membrane (Fig 4A). Experiments with isolated mitochondria further show that while both MPP^+^ and 4′I-MPP^+^ are taken up into isolated MN9D cell mitochondria effectively, the rate of 4′I-MPP^+^ uptake was faster than that of MPP^+^ (Fig 4B). The mitochondrial accumulations of both MPP^+^ and 4′I-MPP^+^ are also associated with a depolarization of the mitochondrial membrane potential (Fig 4C and D) [18]. In parallel to the uptake characteristics, 4′I-MPP^+^ was more effective in depolarizing the mitochondrial membrane potential in comparison to MPP^+^. Specifically, while 500 μM MPP^+^ depolarizes the mitochondrial membrane by 20– 50% in about 45 min, 100 μM 4′I-MPP^+^ takes only 20 min to exert a similar effect on the mitochondrial membrane potential, under similar experimental conditions.

**Fig 4.**
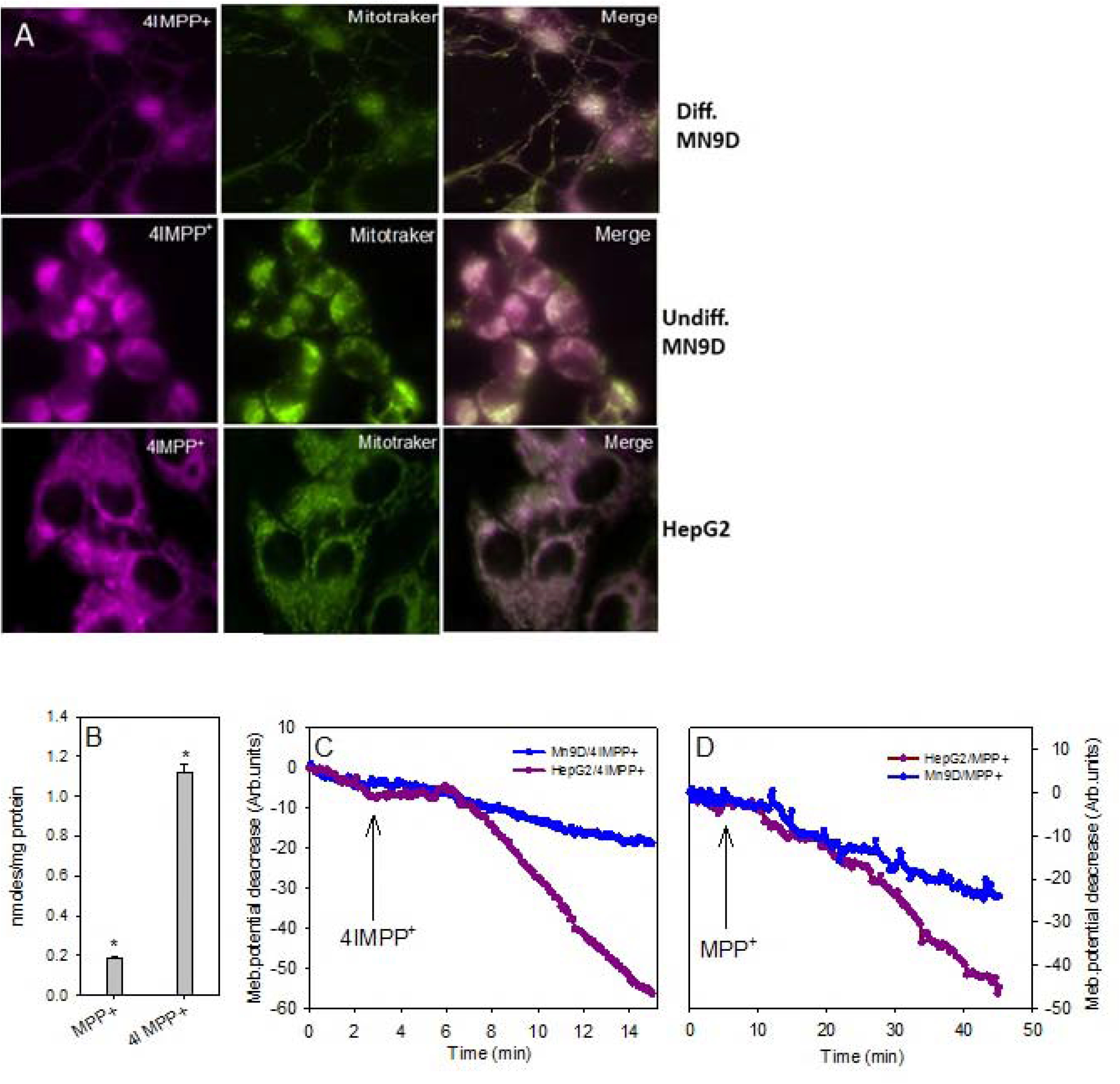
Mitochondrial accumulation and membrane depolarization by 4′I-MPP^+^. (A) Intracellular localization of cytosolic 4′I-MPP^+^. Differentiated, undifferentiated MN9D and HepG2 cells were loaded with 100 μM 4′I-MPP^+^ and 200 nM MitoTracker Green FM, dual fluorescence (4′I-MPP^+^ and MitoTracker Green FM) images were recorded. (B) MPP^+^ and 4′I-MPP^+^ uptake into isolated MN9D cell mitochondria. The uptakes of MPP^+^ or 4′I-MPP^+^ (400 μM; for 45 min at 37 °C) were determined as detailed in Materials and methods. Data presented as mean ± S.D (n = 3). ^*^p< 0.005 versus the zero-time controls; (C and D) **Mitochondrial** membrane potential depolarization by MPP^+^ and 4′I-MPP^+^. MPP^+^ or 4′I-MPP^+^–mediated MN9D or HepG2 cells mitochondrial membrane depolarizations were monitored by observing the decay of intracellular TMRM fluorescence.

### Effect the mitochondrial membrane potential depolarization on the cellular and mitochondrial accumulation of MPP^+^ and 4′I-MPP^+^

The effect of mitochondrial membrane depolarizing agent carbonyl cyanide-ptrifluoromethoxyphenylhydrazone (FCCP) [30, 31] on the cellular accumulations of MPP^+^ and 4′IMPP^+^ were examined. As shown in Fig 5A, 5 μM TFCCP treatment significantly decreased the cellular accumulation of both MPP^+^ and 4′I-MPP^+^ into MN9D cells. Interestingly, the same concentration of FCCP had even a stronger effect on the cellular accumulation of both MPP^+^ and 4′I-MPP^+^ into HepG2, in comparison to MN9D cells. 4′I-MPP^+^ fluorescence imaging experiments show that intracellular 4′I-MPP^+^ in FCCP treated cells is largely distributed throughout the cytosol and not specifically localized in the mitochondria of the cells as opposed to FCCP untreated controls (Fig 5B).

**Fig 5.**
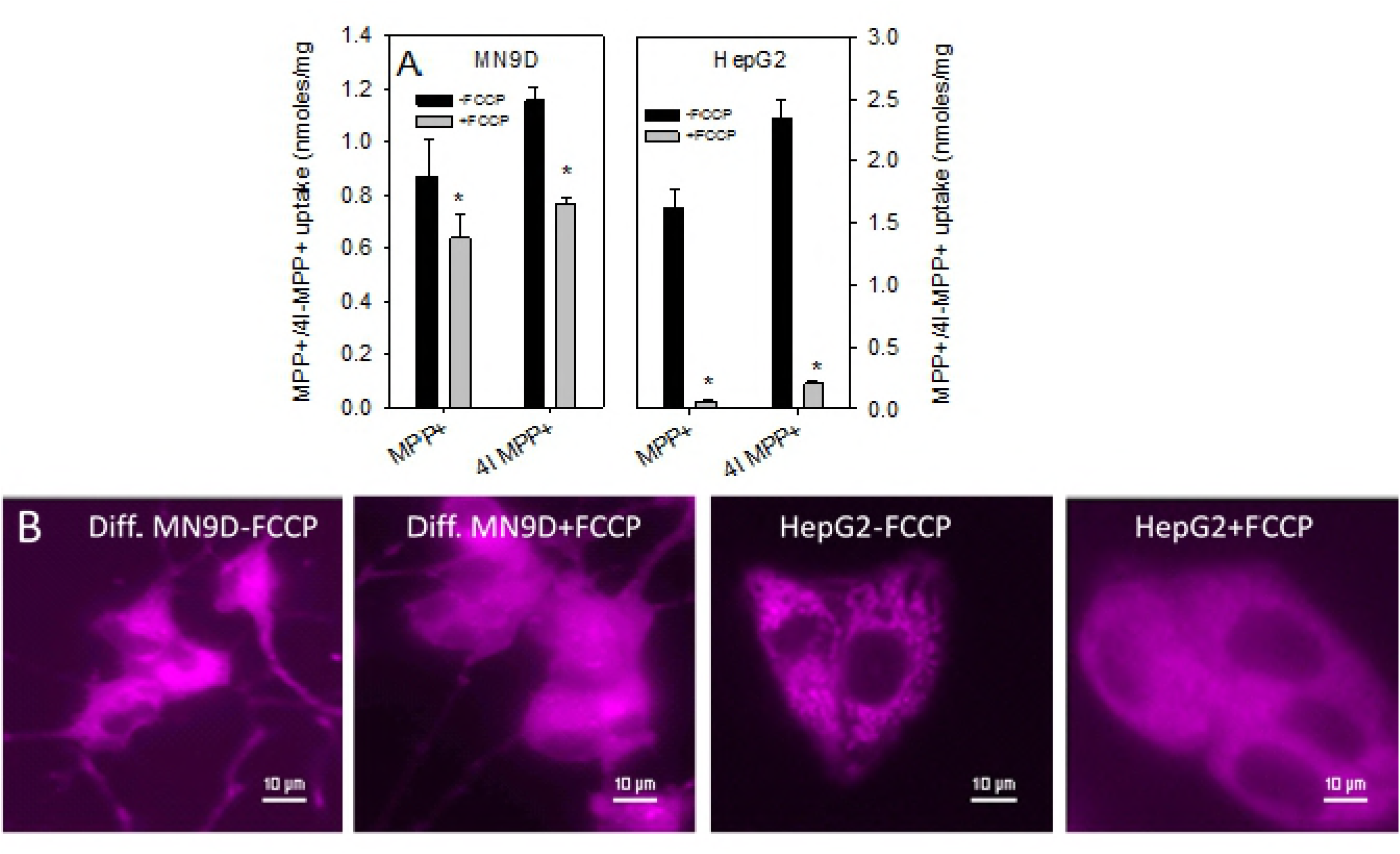
(A) The effect of FCCP on MPP^+^ and 4′I-MPP^+^ uptakes into MN9D and HepG2 cells. Cells were pre-incubated with FCCP (5 μM for 30 min) and then with MPP^+^ or 4′I-MPP^+^ (50 μM, for 45 min at 37 °C) and the intracellular MPP^+^ or 4′I-MPP^+^ levels were quantified as detailed in Materials and methods. The data are presented as mean ± SD (n = 3). ^*^p<0.005 and #p<0.025 versus respective controls without FCCP. (B) The effect of FCCP on the intracellular distribution of 4′IMPP^+^ in differentiated Mn9D and HepG2 cells. Cells were pre-incubated with 5 μM FCCP 30 min and then with 100 μM 4′I-MPP^+^ for 1 h at 37 °C. 4′I-MPP^+^ fluorescence images (Ex/Em 340/470–550 nm) were recorded. Controls were carried out in an identical manner, except that FCCP was omitted from the incubation media.

### Effects of extra-cellular and extra-mitochondrial Ca^2+^ and mNCX and inhibitors on the mitochondrial accumulation of MPP^+^

Previous studies have indicated that intracellular Ca^2+^ may play a role in the cellular [32, 33] and mitochondrial [34] accumulation of MPP^+^. As shown from the data in Fig 6A, 500–1000 nM Ca^2+^ in the incubation media inhibits the 4′I-MPP^+^ uptake into mitochondria by about 40–60% suggesting that extra-mitochondrial Ca^2+^ (i.e. cytosolic) may play a role in the 4′I-MPP^+^ accumulation. In addition, the mNCX inhibitors, CGP37157 [specific mNCX inhibitor[35, 36]], KBR7943[37], and 2APB were also found to inhibit the uptake of 4′I-MPP^+^ (Fig 6B; 50 μM inhibitor at 200 μM 4′I-MPP^+^) into MN9D cells, while the specific mitochondrial Ca^2+^ uniporter inhibitor, ruthenium red [38, 39] (∼10% inhibition was observed at 10 μM concentration), or voltage gated Ca^2+^ channel inhibitors, verapamil [40] or benzamil [41, 42] had no significant effect (S2 Fig). The dose dependency studies with the most effective inhibitor, CGP37157, show that 4′IMPP^+^ uptake inhibition at 10 μM [35] was ∼40% and at 25 μM ∼45% (Fig 6C). To determine whether the mNCX inhibitors are also effective in inhibiting the mitochondrial uptake of 4′I-MPP^+^ in live cells, a series of imaging experiments were carried out with CGP37157. The data presented in Fig 6D show that while cytosolic 4′I-MPP^+^ is primarily localized in the mitochondria (labeled with MitoTracker Green FM) of control HepG2 cells, in cells pre-treated with CGP37157 (50 μM), the intracellular 4′I-MPP^+^ is dispersed throughout the cytosol of the cell. CGP37157 or KBR7943 pre-treatments significantly reduced the typical mitochondrial membrane depolarization seen with 4′I-MPP^+^ in MN9D cells (Fig 6E). The experiments with the mitochondrial membrane potential sensitive fluorescent dye, TMRM, showed that CGP37157 slows down the 4′I-MPP^+^–induced mitochondrial membrane depolarization of HepG2 cells lasting at least 4 h (Fig 6G). The effect of extracellular Ca^2+^ on the uptake of MPP^+^ and 4′I-MPP^+^ into MN9D cells was also examined using standard HPLC-UV techniques. These experiments have revealed that the omission of Ca^2+^ from the incubation medium with the inclusion of a similar concentration of the Ca^2+^ chelator, EGTA, increase the uptake of both toxins by about 3–4 folds (Fig 6F).

**Fig 6.**
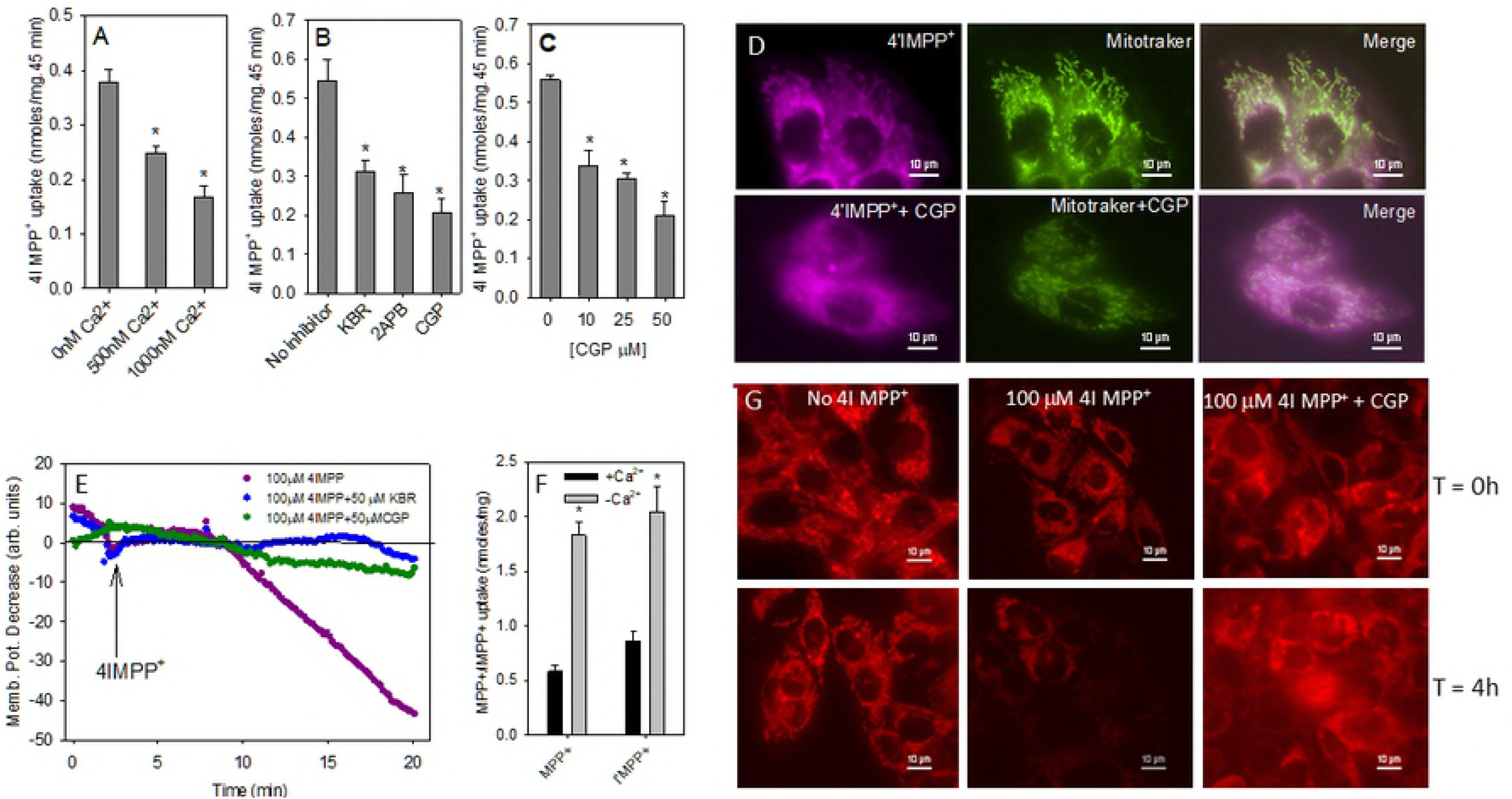
Characteristics of MN9D mitochondrial uptake of 4′I-MPP^+^. (A) Extracellular Ca^2+^ inhibit the mitochondrial uptake of 4′I-MPP^+^. The uptake of 4′I-MPP^+^ (200 μM; for 45 min at 37 °C) into isolated intact MN9D mitochondria were determined as described in Materials and methods in the presence or absence of extracellular Ca^2+^. ^*^p<0.005 versus zero Ca^2+^ controls. (B) Effects of NCX inhibitors on the MN9D mitochondrial uptake of 4′I-MPP^+^. The effects of CGP37157, 2APB, and KBR7943 (50 μM) on the mitochondrial uptake of 4′I-MPP^+^ (200 μM; for 45 min at 37 °C) were determined as detailed in Materials and methods. The data are represented as mean ± S.D (n = 3). *p<0.005 versus controls. (C) Concentration of dependence of the inhibition of MN9D mitochondrial uptake of 4′I-MPP^+^ by CGP37157. The experiments were carried out using a similar protocol as above with varying concentrations of CGP37157 (10–50 μM). The data are represented as mean ± S.D (n = 3). ^*^p<0.005 versus controls. (D) The effect of CGP37157, on the mitochondrial accumulation of 4′I-MPP^+^ in live HepG2 cells. The effect of CGP37157 (50 μM, preincubated for 30 min at 37 °C) on the mitochondrial uptake of 4′I-MPP^+^ (100 μM, for 1 h at 37 °C) was monitored by dual fluorescence imaging using the mitochondrial probe MitoTracker Green and 4′I-MPP^+^. Controls were treated under similar conditions, except that CGP37157 was excluded from the incubation medium. (E) The effect of CGP37157 and KBR7943 on the 4′IMPP^+^–mediated mitochondrial depolarization of HepG2 cells. HepG2 cells were incubated with either 50 μM CGP37157 or KBR7943 for 30 min and then with 50 nM TMRM for 45 min at 37 °C, and finally with 4′I-MPP^+^ (100 μM final concentration). The mitochondrial depolarization is followed by monitoring the intracellular TMRM fluorescence (Ex/Em 543/573 nm) as a function of time. Controls were treated identically except that CGP37157 and KBR7943 were excluded from the initial incubation media. (F) The effect of extracellular Ca^2+^ on MPP^+^ and 4′I-MPP^+^ cellular uptake. Cells were incubated with 50 μM MPP^+^ or 4′I-MPP^+^ in KRB-HEPES or EGTAHEPES (1.3 mM EGTA with no Ca^2+^) for 45 min, at 37 °C. MPP^+^ and 4′I-MPP^+^ uptakes were measured by HPLC-UV. The data are represented as mean ± S.D (n = 3). ^*^p<0.003 versus KRBHEPES controls. (G) The effect of CGP37157 on the 4′I-MPP^+^ mediated mitochondrial membrane depolarization under longer incubation conditions. HepG2 cells were incubated with or without 10 μM CGP37157 followed by 50 nM TMRM, and finally with 4′I-MPP^+^ (100 μM, 0 or 4h) and the intracellular TMRM fluorescence was recorded. Controls were carried out using an identical protocol except that either CGP37157 or 4′I-MPP^+^ were omitted from the incubation media.

### mNCX inhibitors protect MN9D cells from MPP^+^ and 4′I-MPP^+^ toxicities

The inhibition of mitochondrial uptake of both MPP + and 4′I-MPP^+^ suggest that CGP37157 must also protect MN9D cells from the toxicities of both toxins. As shown from the data in Fig 7, CGP37157 pre-treatment effectively protects MN9D cells from MPP^+^ and 4′I-MPP^+^ toxicities, as expected.

**Fig 7.**
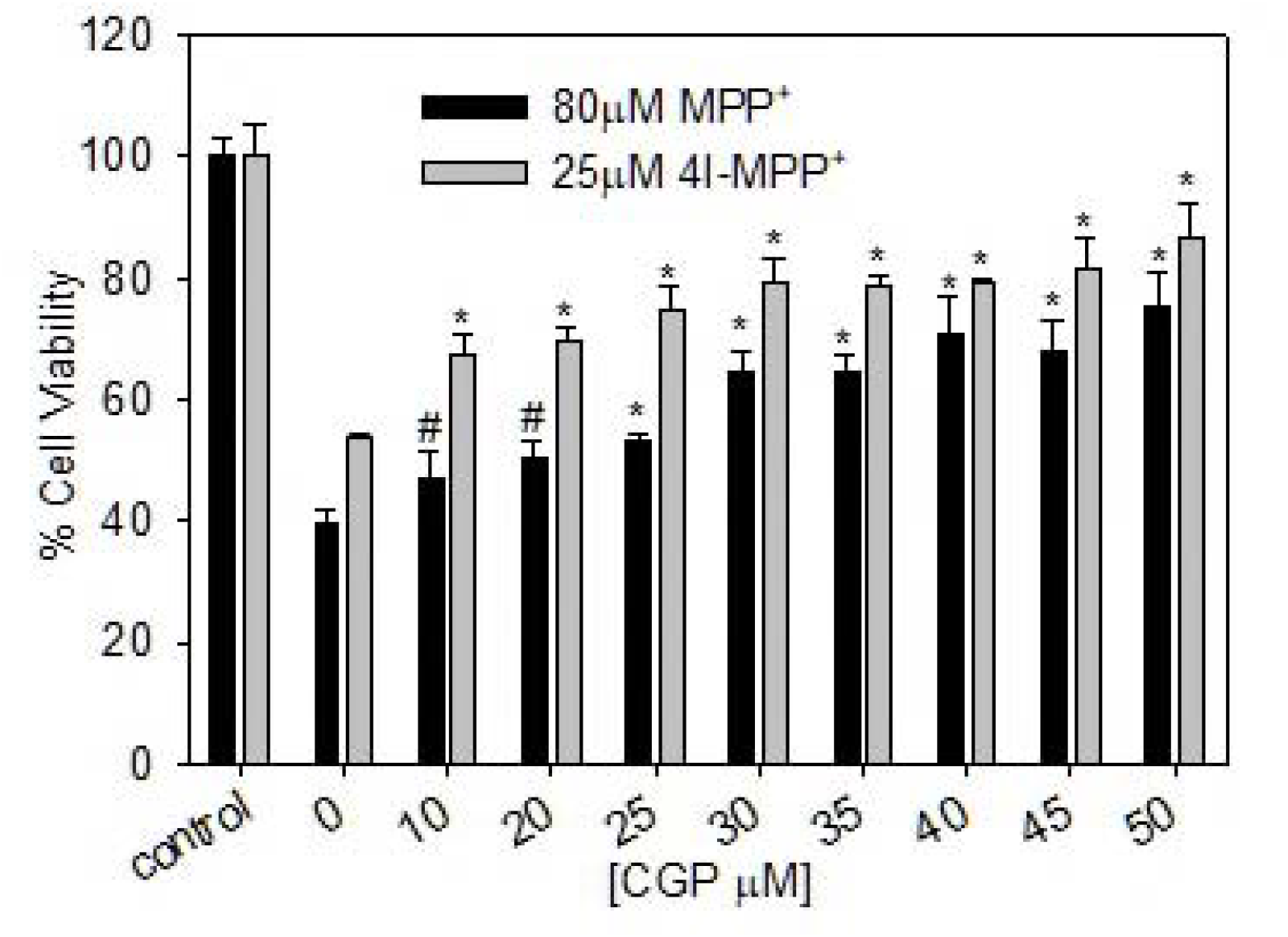
CGP37157 protect MN9D cells from MPP^+^ and 4′I-MPP^+^ toxicities. Cells were pre-incubated with 0–50 µM CGP37157 for 30 min followed by 80 μM MPP^+^ or 25 μM 4′I-MPP^+^ for 8 h. The relative cell viabilities were determiner using MTT assay, as detailed above. Data are presented as mean ± SD (n = 6). ^*^p<0.0001 and #p<0.005 versus no CGP37157 controls.

## Discussion

The aim of this study was to characterize a structural and functional MPP^+^ mimic, which is suitable to study the cellular distribution and mitochondrial uptake of MPP^+^ in live cells and use it to identify the molecular details of these processes to advance the understanding of the mechanism of the selective dopaminergic toxicity of MPP^+^. Accordingly, we have characterized the fluorescent MPP^+^ derivative, 4′I-MPP^+^, as a suitable candidate for this purpose with a number of promising characteristics. 4′I-MPP^+^ which was synthesized in good yield using the previously described procedures,[15] showed a red shifted UV-Vis absorption band at ∼313 nm (with respect to the characteristic UV-Vis band of MPP^+^ at ∼295 nm), and a moderately intense fluorescence band with the excitation and emission maxima at 320 and 430 nm in PBS, respectively. Neither the absorption band nor the fluorescence emission of 4′I-MPP^+^ was significantly blue shifted upon increasing the polarity of the solvent, in contrast to the photophysical behaviors of previously described 4′-(dimethylamino)- and related MPP^+^ derivatives [43, 44]. Similarly, the fluorescence of 4′I-MPP^+^ is moderately quenched by polar solvents, in comparison to 4′-(dimethylamino)-MPP^+^ derivatives and live cell imaging show that intracellular 4′I-MPP^+^ could easily be detected by its intrinsic fluorescence in live calls. These favorable solvatochromic and fluorescent properties of 4′I-MPP^+^ must be associated with its less favorable excited state charge transfer ability in comparison to that of 4′-(dimethylamino)-MPP^+^ derivatives,[45] most likely due to the absence of a highly electron donating 4′-(dimethylamino) functionality in 4′I-MPP^+^. These findings suggest that 4′I-MPP^+^ could be a better behaved fluorescent MPP^+^ mimic which is suitable for in vitro and *in vivo* studies of MPP^+^ toxicity.

Toxicity studies with dopaminergic MN9D [14] cells in reference to non-neuronal liver HepG2 (control) cells [46] have confirmed that 4′I-MPP^+^ is selectively toxic to MN9D cells similar to MPP^+^. In addition, the complex-1 inhibition, cellular ATP depletion, and ROS production selectively in dopaminergic cells leading to apoptotic cell death are also common characteristics of both toxins. However, 4′I-MPP^+^ was a stronger dopaminergic toxin and a complex-I inhibitor in comparison to MPP^+^ under similar conditions, most likely due to its relatively higher cellular uptake and complex–I inhibition potency in comparison to MPP^+^. This notion is consistent with the previously reported relatively high complex-I inhibition potencies and dopaminergic toxicities of 4′-alky substituted lipophilic MPP^+^ derivatives. [47, 48] In addition, the uptake time courses constructed by traditional HPLC-UV and fluorescence measurements under similar conditions were parallel suggesting that the cellular uptake of 4′I-MPP^+^ could be conveniently quantified by fluorescence measurements. Furthermore, the non-specific membrane binding of 4′I-MPP^+^ was insignificant [<1% (data not shown)], in comparison to the more lipophilic 4′-(dimethylamino)-MPP^+^ and related fluorophores, demonstrating that 4′I-MPP^+^ is a convenient structural and functional mimic of MPP^+^ which is suitable to model the cellular uptake, intracellular distribution patterns, and the molecular details of the dopaminergic toxicity of MPP^+^.

Although MPP^+^ is shown to accumulate in the mitochondria isolated from different cell types,[18, 49] the molecular details of the mitochondrial accumulation of MPP^+^ are not fully explored. While the mitochondrial accumulation of MPP^+^ has not been directly demonstrated in live cells, the cellular and various intracellular organelle accumulation of fluorescent 4′-(dimethylamino)-MPP^+^ and related derivatives in live cells have been previously reported [43]. However, the use of these derivatives in the routine cellular or mitochondrial accumulation studies is limited by their nonspecific accumulation into the nucleoli and other organelles (in addition to the mitochondria),[43] interaction and inhibition of various cell membrane transporters, and ability to non-specifically bind to the cell and sub–cellular organelle membranes in large quantities [44,50]. On the other hand, dual mitochondrial and 4′I-MPP^+^ fluorescence imaging show that intracellular 4′I-MPP^+^ is primarily localized into the mitochondria of the cell with only relatively small quantities are associated with the cell cytosol, nucleoli or the membrane (Fig 4A).

Previous studies have suggested that the driving force for the mitochondrial accumulation of MPP^+^ is the large inside negative inner mitochondrial membrane potential [30, 51]. The uptake of MPP^+^ and 4′I-MPP^+^ into isolated MN9D cell mitochondria using traditional HPLC-UV techniques show that the efficiency of 4′I-MPP^+^ accumulation is significantly greater than that of MPP^+^, parallel to the trend that was observed for the cellular uptake. This difference is consistent with the membrane potential facilitated simple diffusion process (*see* below), since 4′I-MPP^+^ is relatively more lipophilic in comparison to MPP^+^ (Table 1). As expected, the mitochondrial accumulation of 4′I-MPP^+^ is associated with a significant depolarization of the mitochondrial membrane potential, as previously reported for MPP^+^ [18]. Furthermore, the more lipophilic 4′IMPP^+^ is a more effective mitochondrial membrane potential depolarizing agent than MPP^+^, which is consistent with the relatively high mitochondrial uptake efficiency of 4′I-MPP^+^ relative to MPP^+^. These findings together with the observation that FCCP (a mitochondrial membrane depolarizing agent) inhibits the mitochondrial accumulation of intracellular 4′I-MPP^+^ in MN9D and HepG2 cells, in comparison to the control cells, suggest the active mitochondrial accumulation of these toxins must be driven by a strong mitochondrial membrane potential energized, simple diffusion process. In addition, a significant decrease of the cellular accumulations of both MPP^+^ and 4′IMPP^+^ in FCCP treated MN9D and HepG2 cells suggest that the effective mitochondrial uptake may also facilitate the cellular uptake of both toxins. Interestingly, HepG2 cell 4′I-MPP^+^ uptake is consistently and significantly more sensitive to FCCP treatments in comparison to MN9D cells. Although the origin of this difference between MN9D and HepG2 cells are not fully understood at present, these findings are in good agreement with a process in which the mitochondrial uptake of MPP^+^ and 4′I-MPP^+^ is a membrane potential facilitated diffusion process in both cell types and the cellular uptakes could be coupled to the mitochondrial uptakes efficiencies at least under *in vitro* conditions.

**Table 1.** The relative elusion times and hydrophobicities of 4′-subtituted MPP + derivatives.

| MPP<sup>+</sup> derivative | Buffer*:CH<sub>3</sub>CN<br>70:30 | Buffer*:CH<sub>3</sub>CN:MeOH<br>60:30:10 | Buffer*:CH<sub>3</sub>CN:MeOH<br>65:25:10 |
| --- | --- | --- | --- |
| 3OH MPP<sup>+</sup> | 5.11 | 4.59 | 5.07 |
| MPP<sup>+</sup> | 5.92 | 5.28 | 6.02 |
| 4F' MPP<sup>+</sup> | 6.44 | 5.68 | 6.60 |
| 4Cl' MPP<sup>+</sup> | 8.66 | 7.08 | 9.17 |
| 4Br' MPP<sup>+</sup> | 9.67 | 7.74 | 10.30 |
| 4I' MPP<sup>+</sup> | 11.76 | 8.88 | 12.56 |

\*Buffer contained 20 mM Na<sub>3</sub>PO<sub>4</sub>, 20 mM CH<sub>3</sub>CO<sub>2</sub>Na, 30 mM triethylamine, 1.7 mM 1-octanesulfonic acid sodium salt, pH 7.0.

Previous studies have indicated that Ca^2+^ may play a role in the cellular[32, 33] and mitochondrial [34] accumulation of MPP^+^. For example, Frei and Richter have reported that Ca^2+^ depleted mitochondria take up more MPP^+^ than Ca^2+^ loaded mitochondria [34]. To model the role of Ca^2+^ in MPP^+^ uptake, the mimic, 4′I-MPP^+^, was used based on the rational that it will produce a clearer picture, due to its relatively high rate of mitochondrial uptake in comparison to MPP^+^.

The isolated MN9D cell mitochondrial 4′I-MPP^+^ uptake is inhibited by extra-mitochondrial Ca^2+^ at physiologically relevant 500–1000 nM concentrations (Fig 6A). More intriguingly, 4′I-MPP^+^ uptake into the mitochondria is also effectively inhibited by mNCX inhibitors CGP37157, KBR7943 and 2APB in a dose dependent manner at pharmacologically relevant concentrations (Fig 6B and 6C). In addition, CGP37157 also prevents the mitochondrial localization of intracellular 4′I-MPP^+^ under the cellular conditions (Fig 6D). In agreement with these findings, CGP37157 also inhibits the characteristic mitochondrial membrane depolarization by 4′I-MPP^+^ in a concentration dependent fashion. These findings suggest that cytosolic/mitochondrial Ca^2+^ play a role in mitochondrial accumulation of these toxins and that this role is likely to be mediated through mNCX. Although, the exact roles of mNCX/Ca^2+^ in cellular/mitochondrial uptake of these toxins are not fully understood at present, repolarization of the toxin depolarized mitochondrial membrane by the efflux of Ca^2+^ through mNCX facilitating the continued electogenic uptake of MPP^+^ or 4IMPP^+^ could be a likely possibility. The observed substantial increase of cellular uptake of both MPP^+^ and 4′I-MPP^+^ into MN9D cells (2 to 3-fold) in Ca^2+^ free, EGTA containing media further suggest that extrusion of the cytosolic Ca^2+^ through plasma membrane NCX may also facilitate the continued uptake of these toxins. These possibilities are in good agreement with the above suggestion that mitochondrial and cellular uptakes of MPP^+^ and 4′I-MPP^+^ are coupled. Since mitochondrial accumulation and the complex-I inhibition mediated increase of ROS production is intimately associated with the MPP^+^ and 4′I-MPP^+^–mediated apoptotic MN9D cell death, [18, 52] the inhibition of mitochondrial uptake by CGP37157 was expected to protect MN9D cells from of these toxins. As expected, CGP37157 effectively protected MN9D cells from both MPP^+^ and 4′IMPP^+^ toxicities (Fig 7). These findings suggest that in addition to mitochondrial accumulation, complex-I inhibition, and ATP depletion MPP^+^ and related mitochondrial toxins may also exert their toxic effects through the perturbation of Ca^2+^ homeostasis in dopaminergic cells. However additional studies are certainly necessary to fully describe the effect of Ca^2+^ on the cellular and mitochondrial uptake and toxicities of MPP^+^ and related toxins. Similarly, to firmly establish the physiological significance of these findings, these *in vitro* cell model finding must be confirmed in physiologically more relevant *in vitro* and *in vivo* model systems.

Taken together, the above findings demonstrate that the unique chemical and photophysical characteristics of 4′I-MPP^+^ together with the its superior uptake and toxicological characteristic relative to MPP^+^ make it a valuable tool to further investigate the mechanism(s) of the selective dopaminergic cell toxicity of MPP^+^ and similar toxins in live cell and animal models. The intrinsic fluorescence of 4′I-MPP^+^ under physiological conditions could also be used to map the cellular distribution of the toxin in the CNS of animal models. The novel finding that cytosolic/mitochondrial Ca^2+^ play a critical role in the mitochondrial and cellular accumulation apparently through NCX is intriguing and suggest for the first time that MPP^+^ and related mitochondrial toxins may also exert their toxic effects through the perturbation of Ca^2+^ homeostasis in dopaminergic cells. Finally, the discovery that specific mNCX inhibitors protect dopaminergic cells from the MPP^+^ toxicity, presumably through the inhibition of the mitochondrial uptake, could potentially be further exploited for the development of pharmacological agents to protect the CNS dopaminergic system from PD-causing environmental toxins.

### Funding Source

Funds to purchase an inverted florescent microscope in part is provided by the National Institutes of Health National Center for Research Resources INBRE Program [Grant P20 RR016475] (to KW).

## Acknowledgement.

We thank Chamila Kadigamuwa for his support in the mitochondrial complex-I inhibition experiments and Shyamali Wimalasena and Nivanthika Wimalasena for critical reading of the manuscript.

## Notes

The authors have no conflict of interest to declare.

## Supporting information

S1 Fig. Standard curves for the RP-HPLC-UV quantification of MPP^+^ and 4′I-MPP^+^

S2 Fig. Effects of Benzamil, Verapamil, FFA, and ruthenium red on the MN9D mitochondrial uptake of 4′I-MPP^+^.

## References

1. Davie CA. A review of Parkinson’s disease. British medical bulletin. 2008;86:109–27. doi: 10.1093/bmb/ldn013. PubMed PMID: 18398010.

2. George JL, Mok S, Moses D, Wilkins S, Bush AI, Cherny RA, et al. Targeting the progression of Parkinson’s disease. Current neuropharmacology. 2009;7(1):9–36. doi: 10.2174/157015909787602814. PubMed PMID: 19721815; PubMed Central PMCID: PMC2724666.

3. Fritz RR, Abell CW, Patel NT, Gessner W, Brossi A. Metabolism of the neurotoxin in MPTP by human liver monoamine oxidase B. FEBS letters. 1985;186(2):224–8. PubMed PMID: 3874094.

4. Nicklas WJ, Vyas I, Heikkila RE. Inhibition of NADH-linked oxidation in brain mitochondria by 1-methyl-4-phenyl-pyridine, a metabolite of the neurotoxin, 1-methyl-4-phenyl-1,2,5,6-tetrahydropyridine. Life sciences. 1985;36(26):2503–8. PubMed PMID: 2861548.

5. Bove J, Prou D, Perier C, Przedborski S. Toxin-induced models of Parkinson’s disease. NeuroRx : the journal of the American Society for Experimental NeuroTherapeutics. 2005;2(3):484–94. doi: 10.1602/neurorx.2.3.484. PubMed PMID: 16389312; PubMed Central PMCID: PMC1144492.

6. Javitch JA, D’Amato RJ, Strittmatter SM, Snyder SH. Parkinsonism-inducing neurotoxin, Nmethyl-4-phenyl-1,2,3,6–tetrahydropyridine: uptake of the metabolite N-methyl-4-phenylpyridine by dopamine neurons explains selective toxicity. Proceedings of the National Academy of Sciences of the United States of America. 1985;82(7):2173–7. PubMed PMID: 3872460; PubMed Central PMCID: PMC397515.

7. Sonsalla PK, Zeevalk GD, German DC. Chronic intraventricular administration of 1-methyl-4-phenylpyridinium as a progressive model of Parkinson’s disease. Parkinsonism & related disorders. 2008;14 Suppl 2:S116–8. doi: 10.1016/j.parkreldis.2008.04.008. PubMed PMID: 18583172; PubMed Central PMCID: PMC2577902.

8. Gainetdinov RR, Fumagalli F, Jones SR, Caron MG. Dopamine transporter is required for in vivo MPTP neurotoxicity: evidence from mice lacking the transporter. Journal of neurochemistry. 1997;69(3):1322–5. PubMed PMID: 9282960.

9. Kalivendi SV, Kotamraju S, Cunningham S, Shang T, Hillard CJ, Kalyanaraman B. 1-Methyl-4-phenylpyridinium (MPP +)-induced apoptosis and mitochondrial oxidant generation: role of transferrin-receptor-dependent iron and hydrogen peroxide. The Biochemical journal. 2003;371(Pt 1):151–64. doi: 10.1042/BJ20021525. PubMed PMID: 12523938; PubMed Central PMCID: PMC1223270.

10. Martel F, Ribeiro L, Calhau C, Azevedo I. Characterization of the efflux of the organic cation MPP + in cultured rat hepatocytes. Eur J Pharmacol. 1999;379(2–3):211–8. Epub 1999/09/25. PubMed PMID: 10497908.

11. Haenisch B, Bonisch H. Interaction of the human plasma membrane monoamine transporter (hPMAT) with antidepressants and antipsychotics. Naunyn Schmiedebergs Arch Pharmacol. 2010;381(1):33–9. Epub 2009/12/17. doi: 10.1007/s00210-009-0479-8. PubMed PMID: 20012264.

12. Ramsay RR, Singer TP. Energy-dependent uptake of N-methyl-4-phenylpyridinium, the neurotoxic metabolite of 1-methyl-4-phenyl-1,2,3,6-tetrahydropyridine, by mitochondria. The Journal of biological chemistry. 1986;261(17):7585–7. PubMed PMID: 3486869.

13. Ramsay RR, Krueger MJ, Youngster SK, Gluck MR, Casida JE, Singer TP. Interaction of 1-methyl-4-phenylpyridinium ion (MPP +) and its analogs with the rotenone/piericidin binding site of NADH dehydrogenase. Journal of neurochemistry. 1991;56(4):1184–90. PubMed PMID: 2002336.

14. Choi HK, Won LA, Kontur PJ, Hammond DN, Fox AP, Wainer BH, et al. Immortalization of embryonic mesencephalic dopaminergic neurons by somatic cell fusion. Brain research. 1991;552(1):67–76. PubMed PMID: 1913182.

15. Wimalasena DS, Perera RP, Heyen BJ, Balasooriya IS, Wimalasena K. Vesicular monoamine transporter substrate/inhibitor activity of MPTP/MPP + derivatives: a structure-activity study. Journal of medicinal chemistry. 2008;51(4):760–8. doi: 10.1021/jm070875p. PubMed PMID: 18220329.

16. Oubrahim H, Stadtman ER, Chock PB. Mitochondria play no roles in Mn (II)-induced apoptosis in HeLa cells. Proceedings of the National Academy of Sciences of the United States of America. 2001;98(17):9505–10. doi: 10.1073/pnas.181319898. PubMed PMID: 11493712; PubMed Central PMCID: PMC55482.

17. Wilkening S, Stahl F, Bader A. Comparison of primary human hepatocytes and hepatoma cell line Hepg2 with regard to their biotransformation properties. Drug metabolism and disposition: the biological fate of chemicals. 2003;31(8):1035–42. doi: 10.1124/dmd.31.8.1035. PubMed PMID: 12867492.

18. Kadigamuwa CC, Le VQ, Wimalasena K. 2, 2’-and 4, 4′-Cyanines are transporter-independent in vitro dopaminergic toxins with the specificity and mechanism of toxicity similar to MPP(+). Journal of neurochemistry. 2015;135(4):755–67. doi: 10.1111/jnc.13201. PubMed PMID: 26094622; PubMed Central PMCID: PMC4793973.

19. Denizot F, Lang R. Rapid colorimetric assay for cell growth and survival. Modifications to the tetrazolium dye procedure giving improved sensitivity and reliability. Journal of immunological methods. 1986;89(2):271–7. PubMed PMID: 3486233.

20. Mosmann T. Rapid colorimetric assay for cellular growth and survival: application to proliferation and cytotoxicity assays. Journal of immunological methods. 1983;65(1–2):55–63. PubMed PMID: 6606682.

21. Zheng XX, Shoffner JM, Voljavec AS, Wallace DC. Evaluation of procedures for assaying oxidative phosphorylation enzyme activities in mitochondrial myopathy muscle biopsies. Biochim Biophys Acta. 1990;1019(1):1–10. PubMed PMID: 2168748.

22. Birch-Machin MA, Turnbull DM. Assaying mitochondrial respiratory complex activity in mitochondria isolated from human cells and tissues. Methods Cell Biol. 2001;65:97–117. PubMed PMID: 11381612.

23. Luo Y, Umegaki H, Wang X, Abe R, Roth GS. Dopamine induces apoptosis through an oxidation-involved SAPK/JNK activation pathway. The Journal of biological chemistry. 1998;273(6):3756–64. PubMed PMID: 9452508.

24. Frezza C, Cipolat S, Scorrano L. Organelle isolation: functional mitochondria from mouse liver, muscle and cultured fibroblasts. Nature protocols. 2007;2(2):287–95. doi: 10.1038/nprot.2006.478. PubMed PMID: 17406588.

25. Joshi DC, Bakowska JC. Determination of mitochondrial membrane potential and reactive oxygen species in live rat cortical neurons. Journal of visualized experiments : JoVE. 2011;(51). doi: 10.3791/2704. PubMed PMID: 21654619; PubMed Central PMCID: PMC3143685.

26. Bradford MM. A rapid and sensitive method for the quantitation of microgram quantities of protein utilizing the principle of protein-dye binding. Anal Biochem. 1976;72:248–54. Epub 1976/05/07. doi: S0003269776699996 [pii]. PubMed PMID: 942051.

27. Przedborski S, Jackson-Lewis V, Naini AB, Jakowec M, Petzinger G, Miller R, et al. The parkinsonian toxin 1-methyl-4-phenyl-1,2,3,6-tetrahydropyridine (MPTP): a technical review of its utility and safety. Journal of neurochemistry. 2001;76(5):1265–74. PubMed PMID: 11238711.

28. Li N, Ragheb K, Lawler G, Sturgis J, Rajwa B, Melendez JA, et al. Mitochondrial complex I inhibitor rotenone induces apoptosis through enhancing mitochondrial reactive oxygen species production. The Journal of biological chemistry. 2003;278(10):8516–25. doi: 10.1074/jbc.M210432200. PubMed PMID: 12496265.

29. LeBel CP, Ischiropoulos H, Bondy SC. Evaluation of the probe 2’,7’-dichlorofluorescin as an indicator of reactive oxygen species formation and oxidative stress. Chemical research in toxicology. 1992;5(2):227–31. PubMed PMID: 1322737.

30. Davey GP, Tipton KF, Murphy MP. Uptake and accumulation of 1-methyl-4-phenylpyridinium by rat liver mitochondria measured using an ion-selective electrode. The Biochemical journal. 1992;288 (Pt 2):439–43. PubMed PMID: 1463448; PubMed Central PMCID: PMC1132030.

31. Tretter L, Chinopoulos C, Adam-Vizi V. Plasma membrane depolarization and disturbed Na + homeostasis induced by the protonophore carbonyl cyanide-p-trifluoromethoxyphenyl-hydrazon in isolated nerve terminals. Molecular pharmacology. 1998;53(4):734–41. PubMed PMID: 9547365.

32. Sheehan JP, Swerdlow RH, Parker WD, Miller SW, Davis RE, Tuttle JB. Altered calcium homeostasis in cells transformed by mitochondria from individuals with Parkinson’s disease. Journal of neurochemistry. 1997;68(3):1221–33 PubMed PMID: 9048769.33.

33. Mohanakumar KP, Ray SS, Büsselberg D. The Parkinsonian neurotoxin 1-methyl-4-phenyl-1,2,3,6-tetrahydropyridine on membrane currents and intrasynaptosomal calcium. Neuroscience Research Communications. 2002;30(1):35–42. doi: 10.1002/nrc.10015.

34. Frei B, Richter C. N-methyl-4-phenylpyridine (MMP +) together with 6-hydroxydopamine or dopamine stimulates Ca2 + release from mitochondria. FEBS letters. 1986;198(1):99–102. PubMed PMID: 3082673.

35. Cox DA, Conforti L, Sperelakis N, Matlib MA. Selectivity of inhibition of Na (+)-Ca2 + exchange of heart mitochondria by benzothiazepine CGP-37157. Journal of cardiovascular pharmacology. 1993;21(4):595–9. PubMed PMID: 7681905.

36. Nicolau SM, de Diego AM, Cortes L, Egea J, Gonzalez JC, Mosquera M, et al. Mitochondrial Na + /Ca2 +–exchanger blocker CGP37157 protects against chromaffin cell death elicited by veratridine. The Journal of pharmacology and experimental therapeutics. 2009;330(3):844–54. doi: 10.1124/jpet.109.154765. PubMed PMID: 19509314.

37. Seki S, Taniguchi M, Takeda H, Nagai M, Taniguchi I, Mochizuki S. Inhibition by KB-r7943 of the reverse mode of the Na + /Ca2 + exchanger reduces Ca2 + overload in ischemic-reperfused rat hearts. Circulation journal : official journal of the Japanese Circulation Society. 2002;66(4):390–6. PubMed PMID: 11954956.

38. Vasington FD, Gazzotti P, Tiozzo R, Carafoli E. The effect of ruthenium red on Ca 2 + transport and respiration in rat liver mitochondria. Biochim Biophys Acta. 1972;256(1):43–54. PubMed PMID: 4257941.

39. Broekemeier KM, Krebsbach RJ, Pfeiffer DR. Inhibition of the mitochondrial Ca2 + uniporter by pure and impure ruthenium red. Molecular and cellular biochemistry. 1994;139(1):33–40. PubMed PMID: 7531818.

40. Davis SE, Bauer EP. L-type voltage-gated calcium channels in the basolateral amygdala are necessary for fear extinction. The Journal of neuroscience : the official journal of the Society for Neuroscience. 2012;32(39):13582–6. doi: 10.1523/JNEUROSCI.0809-12.2012. PubMed PMID: 23015447.

41. Jacob TJ, Civan MM. Role of ion channels in aqueous humor formation. The American journal of physiology. 1996;271(3 Pt 1):C703–20. PubMed PMID: 8843699.

42. Dai XQ, Ramji A, Liu Y, Li Q, Karpinski E, Chen XZ. Inhibition of TRPP3 channel by amiloride and analogs. Molecular pharmacology. 2007;72(6):1576–85. doi: 10.1124/mol.107.037150. PubMed PMID: 17804601.

43. Karpowicz RJ, Jr., Dunn M, Sulzer D, Sames D. APP +, a fluorescent analogue of the neurotoxin MPP +, is a marker of catecholamine neurons in brain tissue, but not a fluorescent false neurotransmitter. ACS chemical neuroscience. 2013;4(5):858–69. doi: 10.1021/cn400038u. PubMed PMID: 23647019; PubMed Central PMCID: PMC3656749.

44. Wilson JN, Ladefoged LK, Babinchak WM, Schiott B. Binding-induced fluorescence of serotonin transporter ligands: A spectroscopic and structural study of 4-(4-(dimethylamino)phenyl)-1-methylpyridinium (APP(+)) and APP(+) analogues. ACS chemical neuroscience. 2014;5(4):296–304. doi: 10.1021/cn400230x. PubMed PMID: 24460204; PubMed Central PMCID: PMC3990943.

45. Ephardt HaF, P. Anilinopyridinium: solvent-dependent fluorescence by intramolecular charge transfer. J Phys Chem. 1991;95(18):6792–7. doi: 10.1021/j100171a011.

46. Kamalian L, Chadwick AE, Bayliss M, French NS, Monshouwer M, Snoeys J, et al. The utility of HepG2 cells to identify direct mitochondrial dysfunction in the absence of cell death. Toxicology in vitro : an international journal published in association with BIBRA. 2015;29(4):732–40. doi: 10.1016/j.tiv.2015.02.011. PubMed PMID: 25746382.

47. Gluck MR, Krueger MJ, Ramsay RR, Sablin SO, Singer TP, Nicklas WJ. Characterization of the inhibitory mechanism of 1-methyl-4-phenylpyridinium and 4-phenylpyridine analogs in inner membrane preparations. J Biol Chem. 1994;269(5):3167–74. Epub 1994/02/04. PubMed PMID: 8106350.

48. Okun JG, Lummen P, Brandt U. Three classes of inhibitors share a common binding domain in mitochondrial complex I (NADH:ubiquinone oxidoreductase). J Biol Chem. 1999;274(5):2625–30. Epub 1999/01/23. PubMed PMID: 9915790.

49. Finichiu PG, James AM, Larsen L, Smith RA, Murphy MP. Mitochondrial accumulation of a lipophilic cation conjugated to an ionisable group depends on membrane potential, pH gradient and pK(a): implications for the design of mitochondrial probes and therapies. Journal of bioenergetics and biomembranes. 2013;45(1–2):165–73. doi: 10.1007/s10863-012-9493-5. PubMed PMID: 23180142.

50. Solis E, Jr., Zdravkovic I, Tomlinson ID, Noskov SY, Rosenthal SJ, De Felice LJ. 4-(4-(dimethylamino)phenyl)-1-methylpyridinium (APP +) is a fluorescent substrate for the human serotonin transporter. The Journal of biological chemistry. 2012;287(12):8852–63. doi: 10.1074/jbc.M111.267757. PubMed PMID: 22291010; PubMed Central PMCID: PMC3308769.

51. Aiuchi T, Shirane Y, Kinemuchi H, Arai Y, Nakaya K, Nakamura Y. Enhancement by tetraphenylboron of inhibition of mitochondrial respiration induced by 1-methyl-4-phenylpyridinium ion (MPP(+)). Neurochemistry international. 1988;12(4):525–31. PubMed PMID: 20501261.

52. Kadigamuwa CC, Mapa MS, Wimalasena K. Lipophilic Cationic Cyanines Are Potent Complex I Inhibitors and Specific in Vitro Dopaminergic Toxins with Mechanistic Similarities to Both Rotenone and MPP(.). Chemical research in toxicology. 2016;29(9):1468–79. doi: 10.1021/acs.chemrestox.6b00138. PubMed PMID: 27510327.

